## Supporting Materials for "Causal Multiscale Modeling of Ocular FLASH Electroretinography: Model Development, Experimental Fitting, and Spin-Glass–Percolation Predictions"

#### Mathematical Derivations, Biophysical Interpretation, and Prospective Predictions


#### S1 Purpose and relationship to the main manuscript

This supporting material is aligned with the author-inclusive Version 8 manuscript used for the arXiv submission. It expands the derivations that are compressed in the article while preserving the same fitted parameter values, nested M0/M1/M2 definitions, and prospective dose-response interpretation. To avoid a notation collision, the normalized optical-stimulus waveform is denoted $\phi_L(t)$ in this supplement, while $s$ is reserved for the Laplace-transform variable and $s_i$ for the Ising diagnostic state. In particular, this supplement provides: (i) the exact solution of the radiation-injury equation; (ii) the Hodgkin–Huxley-to-RC reduction and retinal ion-channel interpretation; (iii) causal derivations of the phototransduction, a-wave, a-to-b, and b-wave stages; (iv) the spin-glass/percolation construction and the meaning of the effective Hamiltonian; (v) the reduced-amplitude benchmark using the Version 8 “red” notation; (vi) a step-by-step derivation of the prospective functional-preservation threshold and dose-modifying factor (DMF); and (vii) additional details on identifiability and experimental validation. The reduced-amplitude fit itself is shown in main-text Fig. 2; the full M2 fit, nested comparison, and network diagnostics are shown in main-text Figs. 3–5.

#### S2 Experimental constraints used by the model

The experimental anchors are the published ocular FLASH electroretinography (ERG) measurements in healthy mouse eyes after single-fraction 21- or 26-Gy irradiation [1]. The reported mean dose rates were $52.6 \pm 2.6$ Gy/s for FLASH and $1.1 \pm 0.1$ Gy/s for conventional-dose-rate (CONV) irradiation, with ERG and histopathologic evaluation at two months. The published study reported five eyes per group. The present calculations use the reported group-mean amplitude-versus-intensity responses as model constraints; accordingly, the fitted parameters should be interpreted primarily as population-level model parameters and prospective hypotheses rather than as uniquely identified microscopic constants.

The X-ray FLASH delivery platform and dosimetry were described previously [2]. The theoretical model deliberately separates the optical light flash that evokes the ERG from the X-ray FLASH irradiation pulse. The optical stimulus enters through $I \to F_L(I) \to L_i(t)$, whereas radiation enters through dose and dose rate, $(D, R) \to X_i(t)$.

#### S3 Dynamic radiation injury

For one coarse-grained retinal module,

$$\frac{dX_i}{dt} + k_{\mathrm{rec,i}} X_i = k_{\mathrm{inj,i}} S_R(R) \dot{D}_i(t). \tag{S1}$$

Multiplication by the integrating factor $e^{k_{\rm rec,i}t}$ gives

$$\frac{d}{dt}\left[e^{k_{\rm rec,i}t}X_i(t)\right] = k_{\rm inj,i}S_R(R)e^{k_{\rm rec,i}t}\dot{D}_i(t). \tag{S2}$$

For a system with no pre-existing model injury in the distant past, integration gives the causal convolution

$$X_i(t) = k_{\rm inj,i}S_R(R)\int_{-\infty}^{t} e^{-k_{\rm rec,i}(t-t')}\dot{D}_i(t')\,dt'. \tag{S3}$$

For a square irradiation pulse of total dose $D_i$ and duration $T_i = D_i/R_i$,

$$\dot{D}_i(t) = \frac{D_i}{T_i}\Theta(t)\Theta(T_i - t). \tag{S4}$$

For $0 \le t \le T_i$,

$$X_i(t) = k_{\rm inj,i}S_R\frac{D_i}{T_i}\int_0^t e^{-k_{\rm rec,i}(t-t')}dt' \tag{S5}$$

$$= k_{\rm inj,i}S_R D_i\frac{1-e^{-k_{\rm rec,i}t}}{k_{\rm rec,i}T_i}. \tag{S6}$$

At the end of irradiation,

$$X_i^*(D_i, R_i) = k_{\rm inj,i}S_R D_i\frac{1-e^{-k_{\rm rec,i}D_i/R_i}}{k_{\rm rec,i}D_i/R_i}. \tag{S7}$$

After the pulse the source term vanishes, so

$$X_i(t) = X_i(T_i)e^{-k_{\rm rec,i}(t-T_i)}, \qquad t > T_i. \tag{S8}$$

The delayed experimental ERG is not assumed to measure the acute endpoint directly. The full fit therefore uses a single global follow-up scale $f_{\rm FU} = 0.487$ to map the acute model injury to the delayed follow-up state. This parsimonious bridge is used because a single follow-up time cannot identify a detailed chronic recovery law.

### S4 Optical light-intensity mapping

The optical light-flash intensity $I$ is mapped to a bounded phototransduction drive through

$$F_L(I) = \frac{I^{n_L}}{I^{n_L} + I_{50}^{n_L}}. \tag{S9}$$

This function satisfies

$$F_L(0) = 0, \qquad F_L(I_{50}) = \frac{1}{2}, \qquad \lim_{I\to\infty} F_L(I) = 1. \tag{S10}$$

The temporal input to module $i$ is

$$L_i(t; I) = L_0 F_L(I)\phi_L(t), \tag{S11}$$

where $\phi_L(t)$ is the normalized *optical* light-flash waveform. It is not the time structure of the X-ray FLASH radiation pulse. Using a common $F_L(I)$ across treatment conditions separates healthy stimulus-response nonlinearity from radiation-induced injury.

#### S5 Hodgkin–Huxley reduction and retinal ion-channel interpretation

The local rate equations are coarse-grained reductions of membrane current balance. For a membrane patch with ionic pathways indexed by $k$,

$$C_m \frac{dV}{dt} = -\sum_k g_k(V,t)\big(V - E_k\big) + I_{\text{drive}}(t), \tag{S12}$$

which has the conductance-balance structure introduced by Hodgkin and Huxley [3]. Linearization around a physiological operating point $V_0$, with the collection of channel-gating variables replaced by an effective slope conductance, gives

$$C_{\text{eff}} \frac{d\delta V}{dt} + G_{\text{eff}} \delta V \simeq I_{\text{drive}}(t), \qquad R_{\text{eff}} = G_{\text{eff}}^{-1}, \qquad \tau = R_{\text{eff}} C_{\text{eff}}. \tag{S13}$$

Thus the $R_a C_a$ and $R_b C_b$ elements in the model summarize effective membrane conductance/capacitance time scales rather than literal electronic components.

The biological pathways represented by the causal states follow the retinal signaling sequence used in the main manuscript. Light activates rhodopsin, transducin, and PDE6, lowers cGMP, and closes cyclic-nucleotide-gated (CNG) $Na^+/Ca^{2+}$ channels. The loss of inward dark current, opposed by $K^+$ efflux, hyperpolarizes photoreceptors [4, 9]. This stage is coarse-grained into $U_i$ and the photoreceptor-dominated state $V_{a,i}$. Photoreceptor hyperpolarization reduces $Ca_V1.4$-dependent ribbon-synaptic glutamate release [5, 9]; the finite signaling state $S_i$ represents this photoreceptor-to-bipolar step. In ON bipolar cells, reduced glutamate decreases mGluR6/$G_o$ signaling, relieves inhibition of TRPM1, and opens a nonselective cation conductance carrying $Na^+$ and $Ca^{2+}$, producing the depolarizing response that contributes strongly to the b-wave [6, 7, 8]. The state $V_{b,i}$ therefore coarse-grains TRPM1-mediated ON-bipolar signaling together with additional $K^+$, $Cl^-$, extracellular, and Müller-cell contributions to the population b-wave.

The present $N = 200$ model does not separately resolve individual Hodgkin–Huxley gating variables, $Na_V$ action-potential generation, OFF-bipolar ionotropic glutamate-receptor pathways, amacrine-cell channel kinetics, or ganglion-cell spiking. This keeps the number of parameters commensurate with the available ERG observables while preserving a direct biophysical interpretation.

#### S6 Laplace transform, inverse Laplace transform, and notation

Because the causal filters below are conveniently solved in the Laplace domain, we define the one-sided Laplace transform of a causal time-domain function $f(t)$ by

$$F(s) \equiv \mathcal{L}\{f(t)\}(s) = \int_0^\infty e^{-st} f(t)\, dt, \qquad s = \sigma + i\omega, \tag{S14}$$

where $t$ is physical time and $s$ is the generally complex Laplace-domain frequency variable. The inverse Laplace transform is

$$f(t) \equiv \mathcal{L}^{-1}\{F(s)\}(t) = \frac{1}{2\pi i} \int_{\gamma - i\infty}^{\gamma + i\infty} e^{st} F(s)\, ds, \tag{S15}$$

where the real constant $\gamma$ is chosen so that the vertical Bromwich contour lies to the right of all singularities of $F(s)$. Two properties used repeatedly below are

$$\mathcal{L}\left\{\frac{df}{dt}\right\} = sF(s) - f(0), \tag{S16}$$

$$\mathcal{L}\{(f * g)(t)\} = F(s)G(s), \qquad (f * g)(t) = \int_0^t f(t - t')g(t')\, dt'. \tag{S17}$$

Thus multiplication of transfer functions in the Laplace domain corresponds to convolution of causal kernels in time. To prevent a notation collision, this supplement reserves $s$ for the Laplace variable and denotes the normalized temporal waveform of the optical ERG stimulus by $\phi_L(t)$. The Ising diagnostic variable $s_i \in \{-1, +1\}$ is separately indexed and should not be confused with either quantity. The main manuscript uses the same $\phi_L(t)$ notation for the optical stimulus waveform, so $s$ remains reserved for the Laplace variable and $s_i$ for the indexed Ising state throughout the arXiv/Overleaf version.

#### S7 Causal phototransduction filter

The first causal state obeys

$$\tau_{r,i}\dot{U}_i + U_i = g_{r,i}L_i(t), \qquad U_i(0) = 0. \tag{S18}$$

Taking a Laplace transform gives

$$(1 + s\tau_{r,i})U_i(s) = g_{r,i}L_i(s), \tag{S19}$$

so the transfer function is

$$H_{r,i}(s) = \frac{U_i(s)}{L_i(s)} = \frac{g_{r,i}}{1 + s\tau_{r,i}}. \tag{S20}$$

For an impulse input, $L_i(t) = \delta(t)$,

$$U_i(t) = \frac{g_{r,i}}{\tau_{r,i}} e^{-t/\tau_{r,i}}\Theta(t). \tag{S21}$$

A finite optical flash is obtained by convolution with the actual light-stimulus waveform $\phi_L(t)$ and produces a causal finite-rise response.

#### S8 Photoreceptor-dominated a-wave and the difference of exponentials

The a-compartment equation is

$$C_{a,i}\dot{V}_{a,i} + \frac{V_{a,i}}{R_{a,i}} = -g_{a,i}U_i, \tag{S22}$$

with $\tau_{a,i} = R_{a,i}C_{a,i}$. For the impulse-driven phototransduction state above and $V_{a,i}(0) = 0$,

$$V_{a,i}(t) = -\frac{g_{a,i}}{C_{a,i}} e^{-t/\tau_{a,i}} \int_0^t e^{t'/\tau_{a,i}} U_i(t')dt' \tag{S23}$$

$$= -A_{a,i}\left(e^{-t/\tau_{a,i}} - e^{-t/\tau_{r,i}}\right), \tag{S24}$$

where $A_{a,i}$ absorbs gain and normalization factors. For $\tau_{r,i} < \tau_{a,i}$, the response starts at zero, develops a finite negative trough, and then recovers. Explicitly,

$$V_{a,i}(0) = 0. \tag{S25}$$

The trough time is obtained from $dV_{a,i}/dt = 0$,

$$t_{a,\text{trough}} = \frac{\tau_{a,i}\tau_{r,i}}{\tau_{a,i} - \tau_{r,i}} \ln\left(\frac{\tau_{a,i}}{\tau_{r,i}}\right). \tag{S26}$$

Thus a-wave timing emerges from causal filtering rather than from an imposed time shift.

#### S9 Explicit a-to-b signaling delay

The downstream signaling state obeys

$$\tau_{ab,i}\dot{S}_i + S_i = -V_{a,i}, \qquad S_i(0) = 0. \tag{S27}$$

Its causal solution is

$$S_i(t) = -\frac{1}{\tau_{ab,i}} \int_0^t e^{-(t-t')/\tau_{ab,i}} V_{a,i}(t')dt'. \tag{S28}$$

Because $S_i$ is a convolution of the already finite-rise a-wave with another causal kernel, it introduces additional processing time and provides a dynamical mechanism for $t_b > t_a$.

#### S10 Post-photoreceptor b-wave and network feedback

The b-compartment satisfies

$$C_{b,i}\dot{V}_{b,i} + \frac{V_{b,i}}{R_{b,i}} = g_{ab,i}S_i + I_{\mathrm{net,i}}, \tag{S29}$$

with $V_{b,i}(0) = 0$ and $\tau_{b,i} = R_{b,i}C_{b,i}$. In the local M0 model, $I_{\mathrm{net,i}} = 0$, giving

$$V_{b,i}(t) = \frac{g_{ab,i}}{C_{b,i}} \int_0^t e^{-(t-t')/\tau_{b,i}} S_i(t')dt'. \tag{S30}$$

The network-augmented models use

$$I_{\mathrm{net,i}}(t) = \eta \sum_j c_{ij} J_{ij} \tanh\left(\frac{V_{b,j}}{V_s}\right). \tag{S31}$$

The hyperbolic tangent is a bounded continuous analogue of the binary Ising variable. In mean-field Ising theory the local average spin has the form $m_i = \tanh(\beta h_i^{\mathrm{eff}})$. Here $\tanh(V_{b,j}/V_s)$ maps a continuous voltage-like state to a signed activity in $[-1, 1]$: for $|V_{b,j}| \gg V_s$ it approaches $\pm 1$, while for $|V_{b,j}| \ll V_s$ it is approximately linear. It therefore provides a soft-spin bridge between continuous ERG dynamics and the binary statistical-mechanical diagnostic while preventing unbounded feedback.

#### S11 Baseline condition and extraction of ERG observables

The initial conditions are

$$U_i(0) = V_{a,i}(0) = S_i(0) = V_{b,i}(0) = 0. \tag{S32}$$

The population ERG is

$$V_{\mathrm{ERG}}(t) = \alpha_a \frac{1}{N} \sum_i V_{a,i}(t) + \alpha_b \frac{1}{N} \sum_i V_{b,i}(t), \tag{S33}$$

so $V_{\mathrm{ERG}}(0) = 0$ exactly.

For optical-flash time $t_{\mathrm{flash}}$,

$$t_a = \arg\min_{t > t_{\mathrm{flash}}} V_{\mathrm{ERG}}(t), \qquad t_b = \arg\max_{t > t_a} V_{\mathrm{ERG}}(t). \tag{S34}$$

For the trough-to-peak convention used in the model,

$$A_a = V_{\mathrm{baseline}} - V_{\mathrm{ERG}}(t_a), \qquad A_b = V_{\mathrm{ERG}}(t_b) - V_{\mathrm{ERG}}(t_a). \tag{S35}$$

The same amplitude convention must be used in model and experiment.

#### S12 Radiation dependence of local electrophysiology

The model permits radiation injury to affect amplitude and timing independently. The representative forms are

$$\tau_{r,i}(X_i) = \tau_{r0,i}(1 + \alpha_{\tau r} X_i), \qquad g_{r,i}(X_i) = g_{r0,i} e^{-\gamma_r X_i}, \tag{S36}$$
$$R_{a,i}(X_i) = R_{a0,i}(1 + \alpha_{R_a} X_i), \qquad C_{a,i}(X_i) = C_{a0,i}(1 + \alpha_{C_a} X_i), \tag{S37}$$
$$R_{b,i}(X_i) = R_{b0,i}(1 + \alpha_{R_b} X_i), \qquad C_{b,i}(X_i) = C_{b0,i}(1 + \alpha_{C_b} X_i), \tag{S38}$$
$$g_{a,i}(X_i) = g_{a0,i} e^{-\gamma_a X_i}, \qquad g_{ab,i}(X_i) = g_{ab0,i} e^{-\gamma_{ab} X_i}, \tag{S39}$$
$$\tau_{ab,i}(X_i) = \tau_{ab0,i}(1 + \alpha_{\tau ab} X_i). \tag{S40}$$

The local baseline quantities define working time and response scales; they should not be interpreted as measured single-channel constants.

#### S13 Spin-glass coupling, coarse-grained binary states, and bond dilution

For a pair of modules,

$$X_{ij} = \frac{X_i + X_j}{2}. \tag{S41}$$

The injury-dependent coupling is

$$J_{ij}(X_{ij}) = \frac{\bar{J}_0}{z_0} e^{-\alpha_J X_{ij}} + \frac{\sigma_0 + \kappa_\sigma X_{ij}}{\sqrt{z_0}} \xi_{ij}, \qquad \xi_{ij} \sim \mathcal{N}(0,1). \tag{S42}$$

The first term describes the coherent contribution and the second describes quenched coupling disorder. Functional connectivity is modeled separately by

$$\Pr(c_{ij} = 1 \mid X_{ij}) = e^{-\alpha_p X_{ij}}. \tag{S43}$$

Here $c_{ij} = 1$ denotes a preserved functional connection and $c_{ij} = 0$ a lost connection. The limiting cases are

$$\alpha_p = 0 \quad \Longrightarrow \quad \Pr(c_{ij} = 1 \mid X_{ij}) = 1, \tag{S44}$$

so there is no radiation-induced bond deletion, and, for any $X_{ij} > 0$,

$$\alpha_p \to \infty \quad \Longrightarrow \quad \Pr(c_{ij} = 1 \mid X_{ij}) \to 0. \tag{S45}$$

At exactly $X_{ij} = 0$, the survival probability remains unity for all $\alpha_p$.

For statistical-mechanical diagnostics each module is assigned

$$s_i \in \{-1, +1\}. \tag{S46}$$

Each $s_i$ represents a coarse-grained local collection of photoreceptors, bipolar cells, interneurons, and supporting retinal cells rather than one neuron or a microscopic magnetic spin. The state $s_i = +1$ denotes a functionally responsive module participating coherently in the stimulus-evoked network response. The state $s_i = -1$ denotes a suppressed, inactive, or oppositely aligned functional state relative to that reference. Because retinal signaling is largely graded, "inactive" does not imply cell death or complete electrical silence.

The diagnostic Hamiltonian is

$$H_{\text{spin}}(X) = -\sum_{i<j} c_{ij}(X) J_{ij}(X) s_i s_j - \sum_i h_i(X) s_i. \tag{S47}$$

Positive $J_{ij}$ favors equal states and negative $J_{ij}$ favors opposite states. A mixture of signed interactions on a surviving network creates competing constraints and frustration [10]. Heat-bath/Glauber updates sample the resulting collective diagnostic states [11]; they do not replace the continuous $U_i \to V_{a,i} \to S_i \to V_{b,i}$ equations that generate the ERG.

The order parameters are

$$m = \frac{1}{N}\sum_i \langle s_i \rangle, \qquad q_{\rm EA} = \frac{1}{N}\sum_i \langle s_i \rangle^2, \qquad q_{\rm SG} = q_{\rm EA} - m^2. \tag{S48}$$

The frustration fraction counts unsatisfied surviving interactions with $J_{ij} s_i s_j < 0$, and the surviving-bond fraction is $p = N_{\rm live}/N_{\rm base}$.

##### S13.1 Ground state and radiation-dependent energy landscape

The Hamiltonian in Eq. (S47) is an effective statistical-physics cost function, not the literal biochemical or electrical energy of retinal tissue. For fixed injury $X$, the ground-state configuration is

$$\{s_i^{(0)}(X)\} = \arg\min_{\{s_i = \pm 1\}} H_{\rm spin}(X). \tag{S49}$$

It represents the collective binary configuration most compatible with the surviving bonds, signed couplings, and local biases. In a healthy-like regime dominated by positive coherent interactions and positive bias, the low-energy state is largely aligned, $s_i \simeq +1$, corresponding to a coherent responsive network. Radiation changes the Hamiltonian itself through $J_{ij}(X)$, $c_{ij}(X)$, and potentially $h_i(X)$,

$$H_{\rm spin}(0) \to H_{\rm spin}(X_1) \to H_{\rm spin}(X_2) \to \cdots. \tag{S50}$$

Thus the relevant picture is a reorganization among the low-energy landscapes of injury-dependent Hamiltonians, not merely excitation of one fixed healthy-eye Hamiltonian. Increased sign disorder and bond loss can create many competing low-energy and metastable states. For the finite $N = 200$ model, changes in order parameters or connectivity should be described as finite-size crossovers or transition regions; a mathematically sharp thermodynamic phase transition would require a large-$N$ analysis and finite-size scaling.

**Why the all-$(-1)$ configuration is not a physical radiation endpoint.** Although the global spin inversion $\{s_i\} \to \{-s_i\}$ is mathematically permissible for an unbiased Ising Hamiltonian, the configuration $s_i = -1$ for every retinal module should not be interpreted as a likely endpoint of radiation-induced evolution from the healthy $s_i = +1$ state. The healthy retina defines a symmetry-selected reference basin, represented in the effective Hamiltonian by the local bias $h_i$, boundary conditions, or conditioning on the initially coherent state. Moreover, the completely inverted state is only one microscopic configuration, whereas radiation-induced disorder makes available an exponentially large number of heterogeneous configurations containing both $+1$ and $-1$ modules. For $N_-$ negative modules the configurational multiplicity is

$$\Omega(N_-) = \binom{N}{N_-}, \qquad S(N_-) = k_B \ln \Omega(N_-), \tag{S51}$$

which is maximal near $N_- \simeq N/2$ but equals zero for the completely ordered limits $N_- = 0$ and $N_- = N$. Thus, particularly in the presence of a symmetry-breaking healthy-state bias, the physically relevant radiation trajectory is from a coherently responsive state toward a heterogeneous ensemble of frustrated, metastable, and potentially disconnected configurations rather than a collective reversal into the unique all-$(-1)$ state. Reduced ERG amplitude is therefore associated primarily with loss of coherent organization and increased configurational disorder rather than with global inversion of all retinal modules.

#### S14 Finite-size percolation diagnostic

Let $n_s$ denote the number of finite connected clusters of size $s$, excluding one largest cluster $S_{\max}$. A standard finite-cluster susceptibility estimator is

$$\chi_{\text{perc}}(p) = \frac{\sum_{s \neq S_{\max}} s^2 n_s}{\sum_{s \neq S_{\max}} s n_s}. \tag{S52}$$

The finite-size pseudocritical point is defined by

$$p_c^{(N)} \simeq \arg\max_p \chi_{\text{perc}}(p). \tag{S53}$$

For the selected $N = 200$, mean-degree-$\simeq 7$ sparse graph ensemble, the independent calculation gives $p_c^{(200)} \simeq 0.17$. This is a topology-dependent model diagnostic, not a universal retinal injury threshold [12, 13, 14].

#### S15 First-order numerical update

Using an explicit Euler step $\Delta t$ and old-time-step values on all right-hand sides,

$$U_i^{n+1} = U_i^n + \frac{\Delta t}{\tau_{r,i}} \left( g_{r,i} L_i^n - U_i^n \right), \tag{S54}$$

$$V_{a,i}^{n+1} = V_{a,i}^n + \frac{\Delta t}{C_{a,i}} \left( -\frac{V_{a,i}^n}{R_{a,i}} - g_{a,i} U_i^n \right), \tag{S55}$$

$$S_i^{n+1} = S_i^n + \frac{\Delta t}{\tau_{ab,i}} \left( -V_{a,i}^n - S_i^n \right), \tag{S56}$$

$$V_{b,i}^{n+1} = V_{b,i}^n + \frac{\Delta t}{C_{b,i}} \left( -\frac{V_{b,i}^n}{R_{b,i}} + g_{ab,i} S_i^n + I_{\text{net,i}}^n \right). \tag{S57}$$

Using the old state on each right-hand side preserves the intended causal ordering within a time step. Higher-order integration may be substituted without changing the model definition.

#### S16 Reduced-amplitude benchmark used in Version 8

The reduced benchmark is intentionally separate from M0. M0 is the local-only realization of the full causal multiscale model; the reduced benchmark is a compact phenomenological amplitude model. Version 8 uses explicit "red" notation to prevent these parameter sets from being conflated.

The reduced injury law is

$$X_{\text{red}}(D, R) = \rho_R \left( \frac{D}{26\,\text{Gy}} \right)^{q_D}, \qquad \rho_{\text{FLASH}} = 1, \qquad \rho_{\text{CONV}} = \rho. \tag{S58}$$

The reduced light-response function is

$$F_L^{\text{red}}(I) = \frac{I^{n_L^{\text{red}}}}{I^{n_L^{\text{red}}} + (I_{50}^{\text{red}})^{n_L^{\text{red}}}}. \tag{S59}$$

For $w \in \{a, b\}$, define the gain-survival factor

$$G_w(D, R) = \exp\left[ -\gamma_w^{\text{red}} X_{\text{red}}(D, R) \right]. \tag{S60}$$

The amplitude prediction is

$$A_{w,c}^{\mathrm{red}}(I;D,R) = \Big[B_{w,c} + A_{w,c}F_L^{\mathrm{red}}(I)\Big]\, G_w(D,R), \tag{S61}$$

so the radiation factor multiplies the complete control response for the corresponding panel.

**Table S1:** Fitted reduced-benchmark parameters used in Version 8.

| Parameter | Value | Interpretation |
|---|---|---|
| $I_{50}^{\mathrm{red}}$ | $0.3459\ \mathrm{cd\,s\,m^{-2}}$ | half-saturation intensity |
| $n_L^{\mathrm{red}}$ | 0.9383 | Hill/Naka–Rushton exponent |
| $\gamma_a^{\mathrm{red}}$ | 0.3751 | a-wave injury sensitivity |
| $\gamma_b^{\mathrm{red}}$ | 0.3300 | b-wave injury sensitivity |
| $\rho$ | 3.3848 | CONV/FLASH effective injury ratio at 26 Gy |
| $q_D$ | 6.4209 | phenomenological dose exponent |
| $B_{a,F}, A_{a,F}$ | 19.304, 51.748 $\mu$V | FLASH-control a-wave floor/range |
| $B_{a,C}, A_{a,C}$ | 23.169, 77.279 $\mu$V | CONV-control a-wave floor/range |
| $B_{b,F}, A_{b,F}$ | 96.226, 122.085 $\mu$V | FLASH-control b-wave floor/range |
| $B_{b,C}, A_{b,C}$ | 126.330, 147.952 $\mu$V | CONV-control b-wave floor/range |

The fitted reduced model gives $R^2 = 0.9721$, RMSE $= 11.94\ \mu$V, and MAE $= 9.36\ \mu$V. From Eqs. (S58) and (S60), the injury scores for FLASH21, FLASH26, CONV21, and CONV26 are

$$X_{\mathrm{red}} = (0.254,\ 1.000,\ 0.859,\ 3.385), \tag{S62}$$

and the corresponding gain-survival factors are

$$G_a = (0.909,\ 0.687,\ 0.725,\ 0.281), \tag{S63}$$
$$G_b = (0.920,\ 0.719,\ 0.753,\ 0.327). \tag{S64}$$

These are derived quantities, not independently fitted parameters.

### S17 Prospective functional-preservation threshold and DMF

Version 8 uses the reduced benchmark to generate a falsifiable prospective dose-escalation prediction. Define $D_{w,r}^*$ as the dose at which the retained gain for ERG component $w \in \{a, b\}$ and irradiation mode $r \in \{\mathrm{FLASH}, \mathrm{CONV}\}$ falls to a prespecified fraction $f_{\mathrm{crit}}$ of control,

$$G_w(D_{w,r}^*, r) = f_{\mathrm{crit}}. \tag{S65}$$

Substituting Eq. (S60),

$$f_{\mathrm{crit}} = \exp\left[-\gamma_w^{\mathrm{red}}\rho_r\left(\frac{D_{w,r}^*}{26\,\mathrm{Gy}}\right)^{q_D}\right]. \tag{S66}$$

Taking the logarithm,

$$-\ln f_{\mathrm{crit}} = \gamma_w^{\mathrm{red}}\rho_r\left(\frac{D_{w,r}^*}{26\,\mathrm{Gy}}\right)^{q_D}, \tag{S67}$$

and therefore

$$D_{w,r}^*(f_{\mathrm{crit}}) = 26\,\mathrm{Gy}\left[\frac{-\ln f_{\mathrm{crit}}}{\gamma_w^{\mathrm{red}}\rho_r}\right]^{1/q_D}. \tag{S68}$$

The functional dose-modifying factor is defined as the FLASH-to-CONV isoeffect-dose ratio,

$$\mathrm{DMF}_w(f_{\mathrm{crit}}) = \frac{D^*_{w,\mathrm{FLASH}}}{D^*_{w,\mathrm{CONV}}}. \tag{S69}$$

Because the same $f_{\mathrm{crit}}$, $q_D$, and $\gamma_w^{\mathrm{red}}$ appear in numerator and denominator, they cancel, yielding

$$\boxed{\mathrm{DMF}_w = \rho^{1/q_D}}. \tag{S70}$$

With $\rho = 3.3848$ and $q_D = 6.4209$,

$$\mathrm{DMF}_{\mathrm{eye}} = 3.3848^{1/6.4209} \simeq 1.209 \simeq 1.21. \tag{S71}$$

Thus the reduced benchmark predicts the same isoeffect ratio for the a- and b-wave under this shared injury law, even though their absolute thresholds differ because $\gamma_a^{\mathrm{red}} \neq \gamma_b^{\mathrm{red}}$.

For the explicitly exploratory choice $f_{\mathrm{crit}} = 0.50$, the thresholds are shown in Table S2 and Fig. S1.

**Table S2:** Prospective 50% retained-gain thresholds derived from the Version 8 reduced benchmark. These are model predictions for experimental testing, not measured retinal-tolerance doses or clinical dose constraints.

| ERG component | $D^*_{\mathrm{FLASH}}$ (Gy) | $D^*_{\mathrm{CONV}}$ (Gy) | DMF |
|---|---|---|---|
| a-wave | 28.6 | 23.7 | 1.21 |
| b-wave | 29.2 | 24.1 | 1.21 |

A conservative combined ERG threshold, defined as the lower of the a- and b-wave thresholds, is therefore approximately 28.6 Gy for FLASH and 23.7 Gy for CONV. The FLASH value is above the highest irradiated calibration dose of 26 Gy and is consequently an extrapolative hypothesis requiring direct dose-escalation validation.

#### S18 Full multiscale fit and nested theory test

For the mechanistic fit, the complete $N = 200$ system is integrated at each treatment condition and optical intensity. A common fixed realization of baseline heterogeneity and quenched graph disorder is retained during optimization so that objective-function changes reflect parameter changes rather than changes in the random realization.

The nested models are

$$\mathrm{M0}: \quad \eta = 0, \tag{S72}$$
$$\mathrm{M1}: \quad \eta \neq 0, \quad \alpha_p = 0, \tag{S73}$$
$$\mathrm{M2}: \quad \eta \neq 0, \quad \alpha_p \neq 0. \tag{S74}$$

Thus M0 contains the complete local causal electrophysiology but no collective network feedback; M1 adds disordered coupling while fixing the topology; and M2 adds both disordered coupling and injury-dependent bond survival.

The scale-balanced objective is

$$\mathcal{L}^* = \sum_{g,k} \left[ \frac{A^{(g)}_{a,\mathrm{mod}}(I_k) - A^{(g)}_{a,\mathrm{exp}}(I_k)}{30\ \mu\mathrm{V}} \right]^2 + \sum_{g,k} \left[ \frac{A^{(g)}_{b,\mathrm{mod}}(I_k) - A^{(g)}_{b,\mathrm{exp}}(I_k)}{65\ \mu\mathrm{V}} \right]^2. \tag{S75}$$

The corresponding nested comparison is summarized in Table S3.

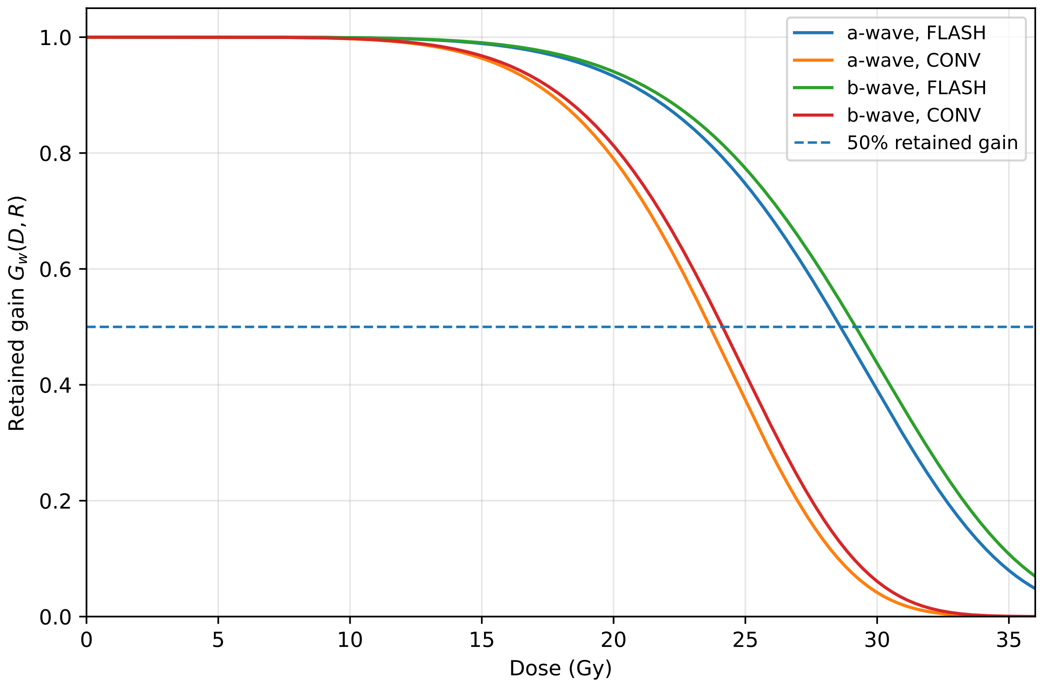


**Figure S1:** Prospective retained-gain curves from the reduced Version 8 benchmark. The dashed horizontal line marks the exploratory $f_{\text{crit}} = 0.50$ criterion. Its intersections with the FLASH and CONV a- and b-wave curves yield the thresholds listed in Table S2. The curves are model predictions, not clinical tolerance curves.

**Table S3:** Nested M0/M1/M2 comparison reported in Version 8. AIC* and BIC* are screening metrics based on the common scale-balanced objective, not a full eye-level likelihood.

| Model | $k$ | $\mathcal{L}^*$ | RMSE ($\mu$V) | $R^2$ | $\Delta$AIC* / $\Delta$BIC* |
|---|---|---|---|---|---|
| M0 | 8 | 25.7721 | 27.2385 | 0.8546 | 0.00 / 0.00 |
| M1 | 11 | 25.6891 | 27.1188 | 0.8559 | 5.69 / 13.38 |
| M2 | 12 | 25.6756 | 27.1059 | 0.8560 | 7.64 / 17.90 |

M2 lowers $\mathcal{L}^*$ by approximately 0.37% relative to M0 and lowers the all-point RMSE by approximately 0.13 $\mu$V. The improvement is small compared with the added model complexity, so the present amplitude measurements do not independently resolve the network terms. This is an identifiability statement, not evidence that collective retinal-network effects are absent.

The profile of the continuous feedback coefficient $\eta$ is shown in Fig. S2. A shallow residual improvement occurs at small network feedback, whereas larger feedback rapidly worsens the fit.

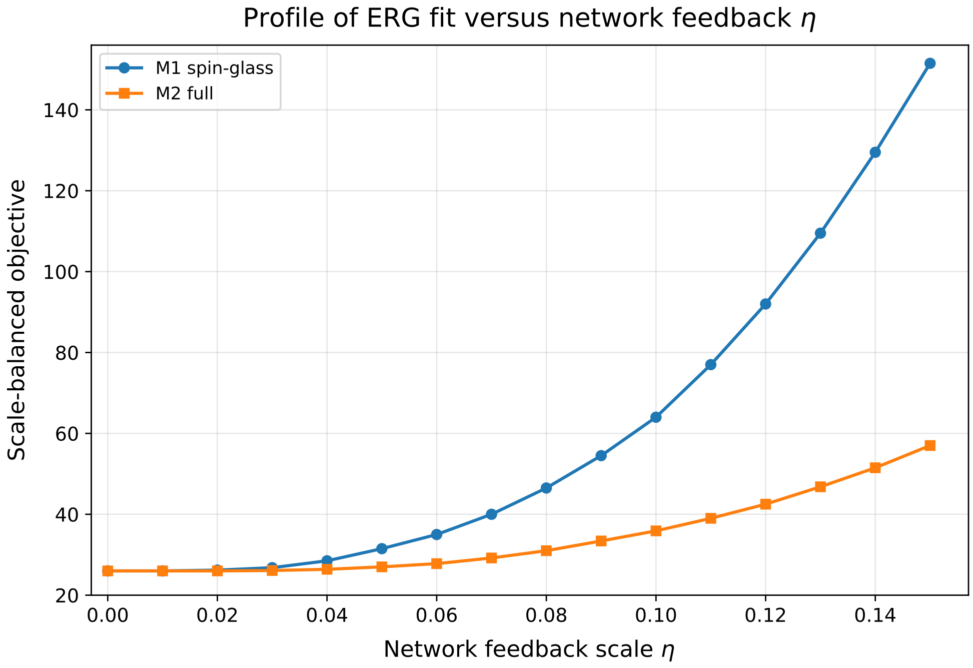


**Figure S2:** Profile of the scale-balanced objective versus b-layer network feedback $\eta$ for M1 and M2. The shallow improvement at small $\eta$ is insufficient to offset the parameter penalty in the nested comparison.

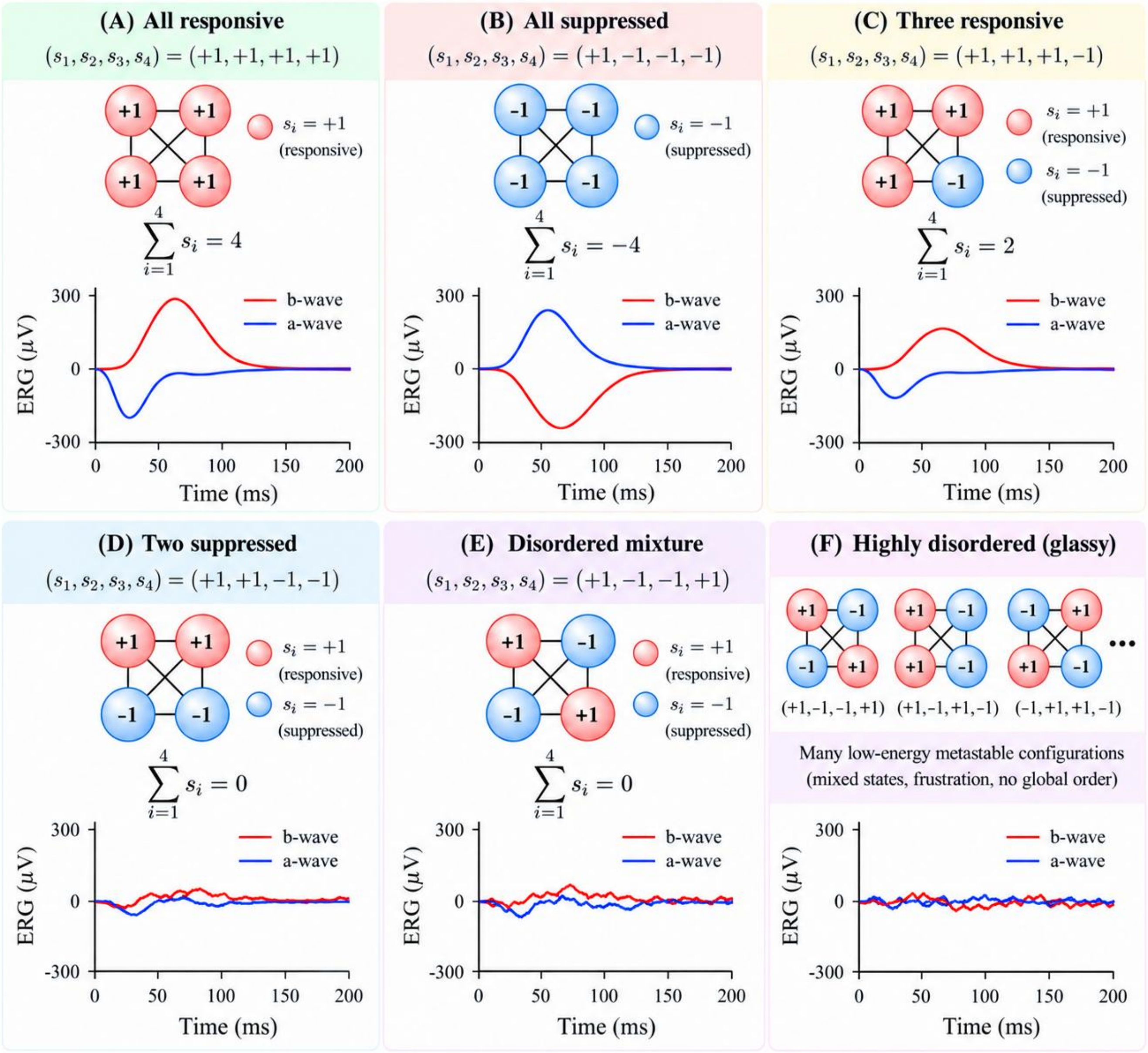


**Figure S3: Illustrative relationship between coarse-grained retinal spin states and ERG output.** Panels A–E show representative $N = 4$ diagnostic spin configurations and schematic population ERG responses, while panel F illustrates the multiplicity of low-energy mixed configurations expected in a glassy regime. The healthy reference state has all modules responsive, $s_i = +1$, and coherent summation produces a large ERG. The globally inverted all-$(-1)$ configuration is shown only as a mathematical comparison state; in the simplified signed-sum illustration it produces the opposite polarity, but it is not interpreted as a likely radiation endpoint. Mixed configurations progressively attenuate or distort the response as coherent population summation is lost. Radiation-induced coupling disorder and frustration are expected to favor an ensemble of competing heterogeneous low-energy or metastable configurations rather than a collective reversal into the unique all-$(-1)$ state. The spin states are coarse-grained functional diagnostics, not literal single-neuron firing states, and the traces are schematic illustrations rather than outputs of the full causal $N = 200$ calculation.

The full M2 fit reported in Version 8 gives $I_{50} = 3.95 \times 10^{-2}$ cd s m$^{-2}$, $n_L = 0.327$, $S_{\text{FLASH}} = 0.507$, and a global follow-up injury scale $f_{\text{FU}} = 0.487$. The effective mean injury scores are approximately 1.03, 1.28, 1.70, and 2.02 for FLASH21, FLASH26, CONV21, and CONV26. The fitted network parameters are $\eta = 0.0141$, $\alpha_J = 0$, $\alpha_p = 0.718$, and $\kappa_\sigma = 1.50$. The boundary tendencies of $\alpha_J$ and $\kappa_\sigma$ indicate limited identifiability and should not be interpreted as direct biological estimates.

#### S19 Why the reduced and multiscale $R^2$ values differ

The reduced benchmark and the multiscale theory test solve different statistical problems. The reduced model is a flexible amplitude-level projection with separate control floor and dynamic-range terms for the two experimental panels and an unweighted amplitude objective. The multiscale fit imposes one causal $N = 200$ architecture, a common model control response, a radiation-injury layer, local causal kinetics, and a nested network hierarchy. Therefore $R^2 = 0.9721$ for the reduced benchmark and $R^2 \simeq 0.856$ for the multiscale fits are not contradictory and should not be compared through a common AIC/BIC calculation. The reduced model calibrates an effective amplitude-response law; M0/M1/M2 tests whether the present observables distinguish local injury from collective network mechanisms.

#### S20 Network predictions and experiments for validation

The fitted M2 realization predicts retained-bond fractions of approximately 1.00, 0.47, 0.39, 0.30, and 0.22 for Control, FLASH21, FLASH26, CONV21, and CONV26, respectively, together with increasing coupling-sign disorder as injury rises. These quantities are model-level predictions rather than directly measured retinal-network observables.

The network theory should therefore be tested with observables that probe collective organization rather than only peak amplitude. Particularly informative measurements include:

1. longitudinal full-waveform ERG across dose, dose rate, light intensity, and follow-up time;
2. repeated-stimulus protocols testing slow relaxation, history dependence, metastability, or hysteresis-like behavior;
3. multifocal ERG, ex vivo multielectrode-array recordings, or calcium imaging for spatial estimates of $m$, $q_{\mathrm{EA}}$, $q_{\mathrm{SG}}$, and pair correlations;
4. dense dose sampling near the predicted preservation region to test changes in fluctuations, susceptibility, or critical slowing;
5. OCT, histopathology, synaptic markers, or related structural measurements to constrain connectivity loss independently of ERG amplitude; and
6. inverse-Ising or maximum-entropy analysis of simultaneous neural recordings to test whether the inferred effective-coupling distribution broadens or develops increased sign disorder after irradiation.

For spatially resolved activity, one may define

$$C_{ij} = \langle s_i s_j \rangle - \langle s_i \rangle \langle s_j \rangle, \tag{S76}$$

and the susceptibility-like quantity

$$\chi = N \left( \langle m^2 \rangle - \langle m \rangle^2 \right). \tag{S77}$$

For connectivity, useful observables include the surviving-bond fraction $p$, largest connected-component fraction $P_\infty = S_{\max}/N$, and finite-cluster susceptibility $\chi_{\mathrm{perc}}$. The strongest validation would combine dose escalation, longitudinal ERG, spatially resolved neural activity, and structural measurements in the same experimental system.

The Version 8 threshold calculation provides a specific near-term design target: measurements should bracket the predicted CONV preservation threshold near 24 Gy and extend FLASH measurements above 26 Gy. The retained-gain criterion should be specified before analysis. A substantially different observed isoeffect ratio would motivate revision of the reduced injury law rather than post hoc redefinition of the threshold.

#### S21 Identifiability and future publication-grade reanalysis

The present fit is based on group-mean amplitude responses. A future eye-level analysis should retain repeated measurements from the same eye and their covariance. A simple hierarchical representation is

$$A_{w,egk} = A^{\mathrm{model}}_{w,gk}(\boldsymbol{\theta}) + b_{w,e} + \epsilon_{w,egk}, \tag{S78}$$

where $b_{w,e}$ is an eye-specific random effect and $\epsilon_{w,egk}$ is measurement noise. Bootstrap resampling should occur at the eye level rather than at individual intensity points.

Full waveforms add information orthogonal to peak amplitudes. With measured timing and waveform uncertainties, one may use a joint objective such as

$$\chi^2_{\mathrm{total}} = \chi^2_{\mathrm{amp}} + \lambda_t \chi^2_{\mathrm{time}} + \lambda_V \chi^2_{\mathrm{waveform}}, \tag{S79}$$

with the weights determined by the measurement covariance or another defensible statistical model. Structural endpoints can be added as independent constraints on bond loss rather than allowing ERG alone to determine both electrophysiology and topology.

#### S22 Reproducibility map

The mechanistic calculation follows

$$(D, \dot{D}) \to X_i(t) \to \{\tau_r, g_r, R_a, C_a, g_a, \tau_{ab}, g_{ab}, R_b, C_b\} \to \{J_{ij}, c_{ij}\} \to V_{\mathrm{ERG}}(t), \tag{S80}$$

while the optical stimulus enters independently through

$$I \to F_L(I) \to L_i(t). \tag{S81}$$

The reduced benchmark instead follows

$$(I, D, R) \to \{F^{\mathrm{red}}_L(I), X_{\mathrm{red}}(D, R), G_w(D, R)\} \to A^{\mathrm{red}}_{w,c}. \tag{S82}$$

The reduced and full calculations are implemented numerically in fixed-form Fortran 77. In the nested multiscale test, the baseline heterogeneity and quenched graph realization are kept fixed during optimization so that differences in the objective reflect model parameters rather than changes of random realization.

#### S23 Summary

The Version 8 framework separates three distinct questions. First, the reduced amplitude benchmark provides a compact description of stimulus-response and radiation-injury dependence. Second, the $N = 200$ causal model tests whether the current amplitude observables distinguish local injury from spin-glass/percolation network mechanisms. Third, the reduced injury law generates a prospective, falsifiable dose-escalation prediction: under the exploratory 50% retained-gain criterion, the conservative combined threshold is approximately 28.6 Gy for FLASH and 23.7 Gy for CONV, with a model-based functional DMF of approximately 1.21. These values are hypotheses for experimental validation, not clinical dose constraints. Current ERG amplitude data support the causal radiation-injury-to-ERG framework, while additional temporal, spatial, correlation-sensitive, structural, and dose-escalation measurements are required to test and distinguish the predicted collective network contributions.
